# Characterisation of cuticle mechanical properties: analysing stiffness in layered living systems to understand surface buckling patterns

**DOI:** 10.1101/2024.03.27.587033

**Authors:** Chiara A. Airoldi, Chao Chen, Humberto Herrera-Ubaldo, Hongbo Fu, Carlos A. Lugo, Alfred J. Crosby, Beverley J. Glover

## Abstract

Development of a living organism is a highly regulated process during which biological materials undergo constant change. *De novo* material synthesis and changes in mechanical properties of materials are key for organ development; however, few studies have attempted to produce quantitative measurements of the mechanical properties of biological materials during growth. Such quantitative analysis is particularly challenging where the material is layered, as is the case for the plant cuticle on top of the plant epidermal cell wall. Here, we focus on *Hibiscus trionum* flower petals, where buckling of the cuticle forms ridges, producing an iridescent effect. This ridge formation is hypothesised to be due to mechanical instability, which directly depends upon the mechanical properties of the individual layers within the epidermal cells. We present measurements of the mechanical properties of the surface layers of petal epidermal cells through atomic force microscopy (AFM) and the uniaxial tensile tester for ultrathin films (TUTTUT), across growth stages. We found that the wavelength of the surface ridges was set at the ridge formation stage, and this wavelength was preserved during further petal development, most likely because of the plasticity of the material. Our findings suggest that temporal changes in biological material properties are key to understanding the development of biological surface patterns.

## INTRODUCTION

*Hibiscus trionum* petals display beautiful colours, which are produced by pigments and structural features. In the purple pigmented regions, the cuticle presents elongated rows of nano-ridges on the top of semi-flat cells, which form a semi-ordered diffraction grating (Whitney et al., 2009). The diffraction of the light by these ridges creates a weak iridescence with a scattered blue halo effect that is perceived by bees and can increase the salience of the flowers to foraging pollinators (Moyroud et al., 2017). The *Hibiscus trionum* petal surface forms a bilayer structure, with the cuticle proper (composed primarily of cutin and waxes) sitting on top of a cuticular layer (composed of cellulose and cutin monomers). It has been shown that mechanical buckling is sufficient to explain the formation of the *Hibiscus trionum* petal cuticular striations (Airoldi et al., 2021).

We have recently explored the theoretical basis of mechanical buckling in the *Hibiscus trionum* system (Lugo et al., 2023). According to linear elastic buckling theory, a bilayer system with a stiff thin film resting on a soft substrate will form a periodic wrinkle pattern when an in-plane compressive strain exceeds a critical threshold value, *ε*_*c*_ (Bowden et al., 1998; Chan and Crosby, 2006; Cai et al., 2011; Efimenko et al., 2005; Huck et al., 2000). If the substrate thickness is much larger than the film thickness, the wavelength at the onset of a wrinkled pattern in an incompressible material *λ* is determined by:

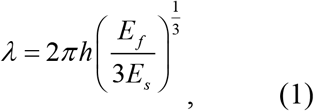

Chen et al., 2020), where *E*_f_ and *E*_s_ are the Young’s moduli of the film and the substrate, and *h* is the film thickness. The Young’s modulus (E) is a property of the material that tells us how easily it can stretch and deform and is defined as the ratio of tensile stress (σ) to tensile strain (ε), where stress is the amount of force applied per unit area (σ = F/A) and strain is extension per unit length (ε = dl/l). The higher the modulus, the more stress is needed to create the same amount of strain.

Mechanical buckling deformation mechanisms have been shown to describe the wrinkle patterns on *Hibiscus trionum* and to fit with known descriptions of cell and tissue morphology (Lugo et al., 2023). However, quantitative comparisons are limited since they require precise knowledge of the mechanical properties of the film and substrate. The formation and morphology of such patterns are closely related to the mechanical properties of the constitutive materials and determined by the ratio E_f_/E_s_. In addition, knowledge of how the mechanical properties of the materials change during growth would aid our understanding of how specific wrinkle patterns and dimensions are achieved as the flower petals mature. While it seems intuitively likely that the mechanical properties of biological materials change during development, very few studies have been dedicated to measuring such changes in living, growing tissues.

In this work, we developed tools to measure the mechanical properties of the cuticle proper and the cuticular layer in the petals from the onset of buckling to the fully developed flower. We used two independent methods, atomic force microscopy (AFM) and the uniaxial tensile tester for ultrathin films (TUTTUT), to measure the elastic moduli of the constitutive layers of the petal surface and assess the reproducibility of our data. By directly characterizing the material and geometric properties, we were able to assess whether the morphology we observe can be predicted by the mechanical theory. We found that the properties of the tissue at the start of buckling can predict the wrinkling initiation strain and the wrinkle wavelength. We also found that the pattern wavelength is set immediately following initiation, such that the later development of the petal does not substantially change the pattern wavelength. This finding indicates a deviation from the wrinkle pattern prediction using mature stage properties, and the likelihood that the locking of the pattern wavelength can be attributed to the inelasticity of biological materials during growth. The measurement of mechanical properties during development will provide new insights in the field of biomechanics and the development of bioinspired materials systems.

## MATERIALS AND METHODS

### Plant growth conditions

*Hibiscus trionum* seeds were obtained from Cambridge University Botanic Garden. Plants were grown in Levington’s M3 compost in a greenhouse with a controlled temperature of 21 ºC, and a 16-hour daylight regime. Buds were measured using a Vernier caliper after removing the epicalyx and measuring from the base of the bud to the top.

### Bright field light microscopy

A VHX-5000 Keyence microscope with a VH-Z500R/W/T 392 objective was used for bright field microscopy.

### Atomic Force Microscopy (AFM)

An AFM JPK NanoWizard with a cantilever tip of tetrahedral shape (ATEC-CONT Au10 S/N: 74454F6L1180 Nanosensor; SUPPLEMENTAL FIGURE S01) was used. This cantilever was chosen because of its small half cone angle that increases its performance on samples with a small pattern size combined with steep sample features. The cantilever’s spring constant was calibrated using the thermal noise method in the air. Three cantilevers were used, all with measured spring constants within the manufacturer’s stated range (0.25 N/m, 0.16 N/m and 0.26 N/m). The sensitivity was calculated in water at the beginning of every set of measurements by fitting the slope of a force-displacement curve acquired on a stiff substrate (a glass slide). The measured values ranged from 50 to 80 nm/V. We used the JPK software (JPK00806) in force mapping mode. Force-displacement curves were recorded within an area of 1 um x 1 um with a grid of 32 x 32 points. Force–displacement curves were recorded at a speed of 40 um/s.

Samples were prepared by positioning a small square of the peeled epidermis on a glass slide surrounded by 1 % low melting point agarose. Alternatively, we positioned the samples on a cooled but not completely solidified 3 % agarose gel. We did not observe differences in our measurements between the two methods, but we observed that the 3 % agarose was more reliable to avoid movement of the samples. Measurements were conducted in water with a small amount of sucrose (0.005 M). We measured the cuticular layer (substrate) following removal of the cuticle proper (upper layer) using a sharp and flexible blade while monitoring the process under a stereomicroscope.

The JPK Imaging mode was used to acquire the image of the peeled flower. The area scanned was 20 um x 20 um with a grid of 128 x 128 pixels. The tip velocity was 13.4 um/s, and a set point of 5 nN was used. The depth of the indentations was never more than 1/10 of the layer measured, and this was repeatedly monitored during indentation by processing curves – any measurements to greater depth were excluded from the analyses. For the cuticle at stage 3, we used a cut-off value of 29 nm. For the cuticle at stage 5, the cut-off value was 64 nm. The imaging was exported directly from the cantilever height measured without a plane or line subtraction to give a better idea of the overall shape of the petal surface. Measurements were made on petals from at least three independent flowers per treatment, analysing multiple regions within a petal, and three measurements per region were performed, total measurement numbers for each stage are indicated in Figure 3.

JPK SPM Data Processing BRUKER software was used for data analysis. Baseline subtraction (to subtract for the baseline offset), contact point (to adjust the X offset) and vertical tip position (to subtract the cantilever deflection in the unit of length from the piezo height, to correct the height for the bending of the cantilever to calculate the vertical tip position) were all employed. Elasticity fit was calculated on the extended curve to avoid artefacts given by adhesion present in the retract curve. Zemla et al. (Zemla et al., 2020) showed that pyramidal shapes fit for a pyramidal tip do not give accurate Young’s modulus values. The choice of contact mechanics model was determined considering the scaling of the measured force-displacement relationships, as well as microscopy analysis of the SPM tips, which indicated that the contact area between the sample and the tip did not change significantly during unloading. The Hertz model, based on the contact of a flat cylindrical tip with a constant radius (radius = 10 nm), was applied to extract Young’s moduli values.

### TUTTUT

The cuticle proper was separated from *H. trionum* petals using an enzymatic protocol modified from a previous method of cuticle isolation from tomatoes (España et al., 2014; Petracek et al., 1995). In brief, petals were treated in an aqueous enzyme bath consisting of fungal cellulase (0.2 % w/v), pectinase (2.0 % w/v), and sodium azide (1 mM) in sodium citrate buffer (pH 3.7, Millipore Sigma Corporate). Petals were placed on a glass plate, and the abaxial side of the petals was scratched off with a razor blade such that the components beneath the adaxial cuticle were exposed for enzymatic treatment. The petal was submerged in the enzyme solution in an enclosed container to avoid evaporation and incubated at 35 °C for 24 hours. The isolated cuticle was then moved using tweezers to a glass slide and rinsed with deionized water. We used an optical profilometer to measure the thickness of the individual cuticles.

We followed the protocol described in Bay et. al, 2019 to conduct the The Uniaxial Tensile Tester for UltraThin films (TUTTUT) to measure the force–displacement responses (Bay et al., 2019). The TUTTUT cantilevers were calibrated before each set of measurements, following protocols previously published (Bay et al., 2019). We floated the cuticle films on a water bath. A silicon wafer was dropped on the grip section of the film. The water level was lowered, and the wafer positioned into the clamp and rigidly fixed onto the walls of bath container. The other side of the cuticle film was attached to a cantilever (an aluminium-coated cover glass). The grip section wafer was set to be parallel to the bottom edge of the cantilever. The stage connected to the water bath container was then pulled at a constant velocity of 0.2mm/s. A laser reflective system recorded the cantilever deformation, which was then used to calculate the resulting force on the cuticle film. For each petal developmental stage, we analysed three isolated cuticle films. We used the force–displacement responses to determine the average stress–strain responses of the cuticle samples. The strain was calculated by dividing the applied far-field displacement by the initial sample length. Since the shapes of the isolated cuticle samples were irregular (i.e., not rectangular nor standard geometries), we used finite element analysis (FEA) with the same geometries as the experiments to calibrate a nominal cross-sectional area in the loading direction for each experiment to determine the Young’s modulus of the cuticles. Specifically, the Young’s modulus of the FEA model was adjusted until the relationship between resultant force and applied displacement matched experimentally measured trends for each sample geometry. We then obtained the nominal cross-sectional area in the loading direction by dividing the experimentally measured stiffness by the determined Young’s modulus. The stress was then determined by dividing the resultant force by the nominal cross-sectional area.

The finite element calibrations of the TUTTUT experiments to obtain the nominal cross-section area were conducted using linear elastic plane stress analyses in ABAQUS software. ABAQUS plane stress elements CPS4 (dominating element) and CPS3 were used. A typical analysis was shown in Figure S5-B-C for a cuticle sample at the striation onset stage (Figure S5-A). The mesh sensitivity was checked by varying mesh size from 2mm to 0.02mm, with negligibly small variation of 0.57% of the resulting force output as well as the nominal area (Figure S5-D). No remeshing was conducted for the small strain analysis.

## RESULTS

### 1. Characterization of growth of *Hibiscus trionum* petal layers

Petal development in *H. trionum* involves changes in size, shape, colour, and texture. The cuticular striation patterns in the purple-coloured section of *H. trionum* petals take around 3 days to develop, immediately before flower opening. During this time the flower bud also increases its size dramatically (Moyroud et al., 2022). FIGURE 1A-D summarizes cuticular striation development as a function of time. In three days, a bud (less than 10 mm), develops from a pre-striation stage displaying smooth cuticles, to an onset stage, then to a mature bud (around 13 mm bud size), which displays ordered striations, and finally to a fully developed and open flower. Note that the white region of the petal is composed of epidermal cells with a conical shape and a smooth cuticle that does not form striations (Moyroud et al., 2022).

**Figure 1.**
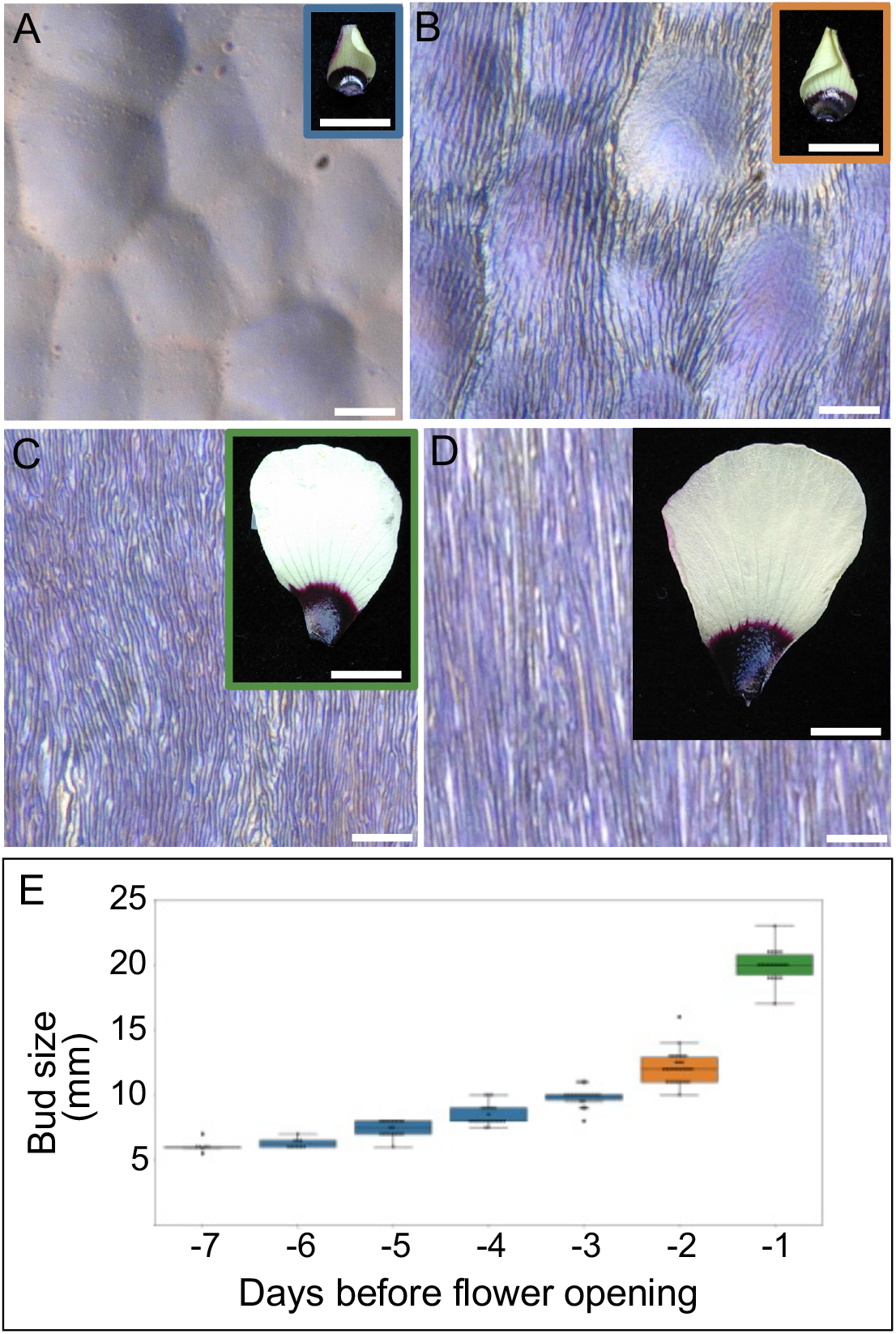
Petal growth and striation formation in *Hibiscus trionum*. (A) A 10 mm flower bud with smooth petal epidermis (pre-striation stage). (B) 12mm flower bud petal with striations starting to form. (C) Mature flower bud petal with striations formed across the purple area of the petal. (D) Open flower petal with fully developed semi-ordered striations. (E) Flower bud size (without epicalyx) from 7 days before opening. Pre-striation stages (-7 to -3 days) are indicated in blue bars, the striation onset stage (-2 days) in orange, and the mature bud stage in green; these colours also indicate stage around the petal images in A-D. Scale bars represent 10 μm in A-D, and 10 mm in the insets.

We performed a detailed characterization of the thickness of the substrate, the cuticular layer, prior to mechanical measurements. We have previously measured the thickness of the cuticle proper (the upper layer) at different stages of petal development from cryoSEM fracture images of the entire layered structure, taking measurements from both the crest and the base of the ridges (Lugo et al., 2023, also summarized in TABLE 1).

**Table 1.** Summary of cellular features and cuticle material properties used to calculate the wavelength values using equation 1. in Lugo et al., 2023 are marked with *.

| Sample | Thickness |  |  | E, Young modulus (AFM) |  | Wavelength |  |
| --- | --- | --- | --- | --- | --- | --- | --- |
|  | Cuticle proper*<br>nm (st dev) | Cut. layer top*<br>nm | Cut. layer valley*<br>nm | Cuticle proper<br>MPa (st dev) | Cuticular layer<br>MPa (st dev) | Theoretical<br>nm | Observed*<br>nm |
| Pre-striation | 89.0 | 1790.8 | 1790.8 |  |  |  |  |
| Onset | 146.3 (25.7) | 2544.2 | 2429.9 | 27.3 (9.57) | 7.3 (13.48) | 987 (±646.27) | 1308.8 |
| Mature bud | 210.0 | 2605.0 | 2171.6 | 24.8 | 1.2 | 2510.0 | 1289.3 |
| Open flower | 193.0 | 1879.9 | 1473.4 | 20.1 | 1.1 | 2115.0 | 1313.8 |

Here, to measure the thickness variation of the cuticle proper across the entire petal surface, we used an optical profilometer (Zygo, Nextview NX) with our isolated cuticle preparations (FIGURE 2). The thickness of the isolated cuticle proper varied from 150 nm to 350 nm across the petal at the open flower stage (FIGURE 2E), similar to the range of measurements recorded from non-isolated layered images (TABLE 1). The cuticle proper thickness is around 0.2 to 0.4 um across the petal, with some peaks that can reach 0.8 um, additionally, the thickness in the central region is higher (FIGURE 2E). We therefore focus on the central part of the purple region of the petal, where striations develop.

**Figure 2.**
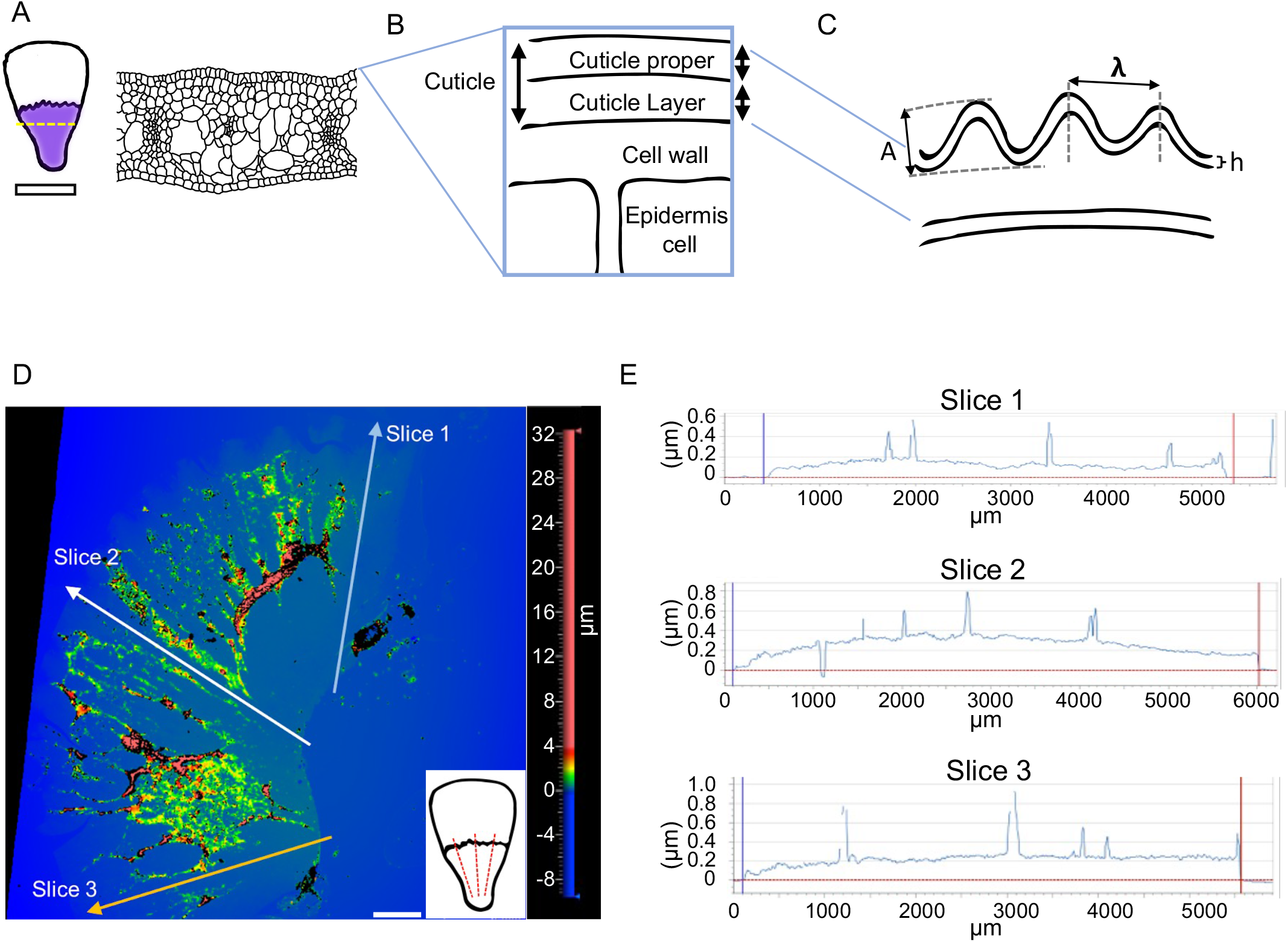
Cuticle thickness analyses. (A) Cuticle striations are formed on the epidermal cells in the purple region in the petals. (B) Schematic representation of cuticle and its parts: cuticle proper (cutin + wax) and cuticle layer (cutin + wax + pectins). (C) Features of the wrinkles, A, amplitude; λ, wavelength; h, film thickness. (D) Profilometer analysis of cuticle proper removed from the petal. (E) Thickness was measured in 3 slices across the petal represented by the red dashed lines in the inset in (D). Scale bar represents 1 mm in D.

The cuticular layer increases in thickness during striation onset and through early striation development, (TABLE 1), then it shows a slight reduction in thickness as the flower surface grows considerably at the final stage of bud development (FIGURE 1E, SUPPLEMENTAL FIGURE S02). This observation is consistent with rapid cell surface growth and mass increase during this stage.

In summary, the growth of the *H. trionum* petal involves changes in the thickness of both layers of the cuticular bilayer that forms the striations.

### 2. Characterization of Young’s moduli of the cuticle proper and the cuticular layer

A number of papers have suggested that the cuticular striations of *H. trionum* develop as a result of mechanical instability (Antoniou-Kourounioti et al., 2012; Chen et al., 2020), with our latest model (Lugo et al., 2023) proposing that their formation is the result of differences in the mechanical properties of two layers within the cuticle: the cuticle proper (top layer) and the cuticular layer (bottom layer). The model tested a range of values of stiffness ratio between the two layers and defined a value for the formation of striation patterns.

In this work, we assumed the cuticle is an isotropic material, so, measurements on the z-component could give us an idea about the in-plane properties. We used Atomic Force Microscopy (AFM) and The Uniaxial Tensile Tester for UltraThin films (TUTTUT) to measure Young’s moduli of the cuticle proper and we used AFM to take similar measurements from the cuticular layer (FIGURE 3). FIGURE 3B shows the appearance of the petal after removal of the cuticle proper, revealing the cuticular layer. FIGURE 3C shows the appearance of an isolated cuticle proper. The results of our TUTTUT experiments to analyse the cuticle proper are summarized in FIGURE 3D,E for striation onset and open flower stages. The mean Young’s modulus of the cuticular layer at striation onset is around 7.3 MPa, decreasing to around 1.2 MPa at the open flower stage. This decrease in stiffness of the cuticular layer during tissue development has not previously been described.

**Figure 3.**
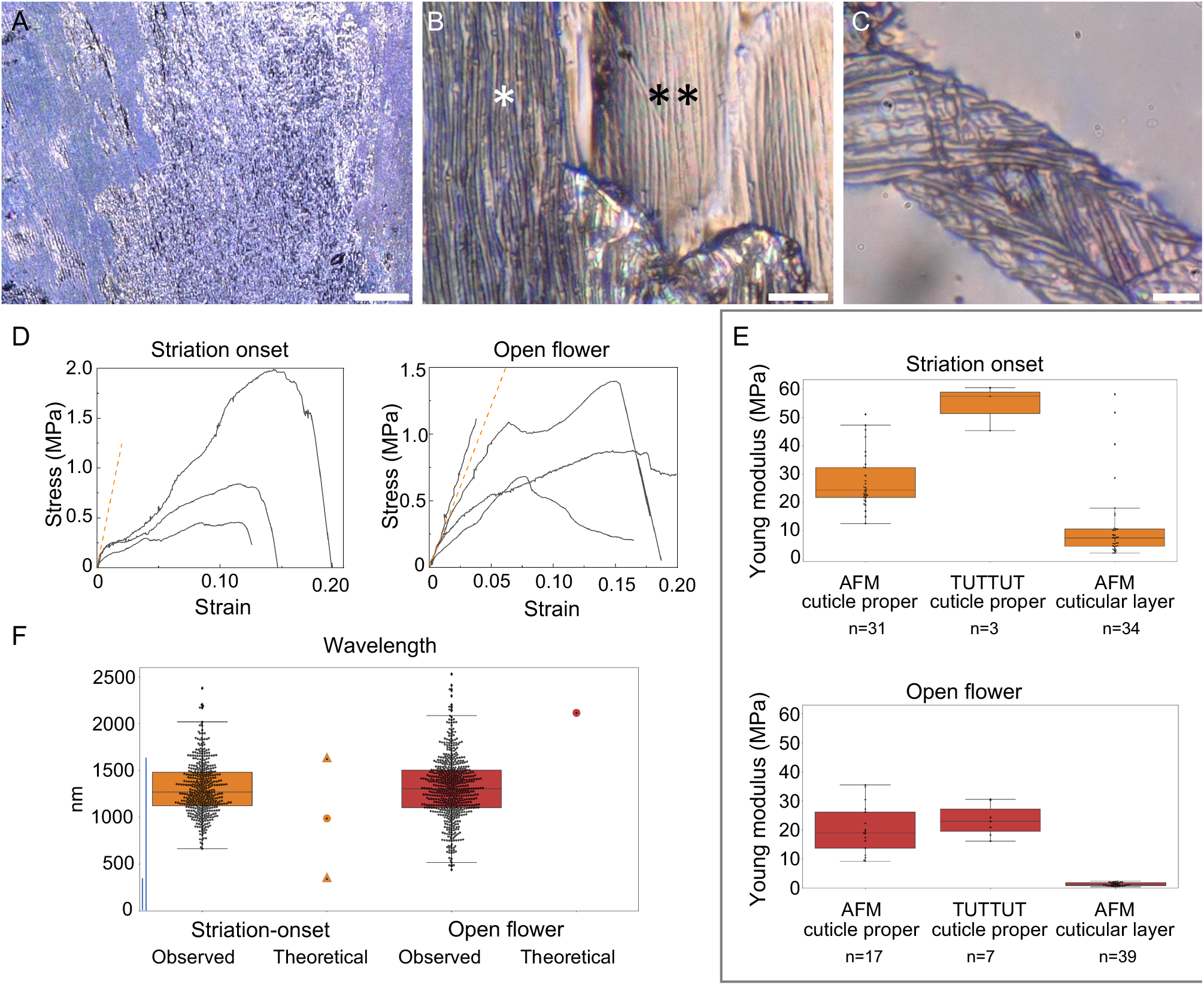
Characterization of cuticular layers stiffness during striation in *Hibiscus trionum* petals. (A,B) Keyence images of a petal before and after cuticle proper removal. (B), Overview of a petal region after cuticle proper removal, cells still with the cuticle proper are indicated with *, cells without the cuticle proper are marked with **. (C) Isolated cuticle proper after being removed from the petal. (D) Stress vs strain values from the uniaxial tensile tester for ultrathin films (TUTTUT) measurements of the cuticle proper at the striation onset and at the open flower stage. (E) Young’s moduli values calculated from Atomic Force Microscopy (AFM) and the TUTTUT measurements during striation onset and at the open flower stage. (F) Observed vs theoretical wavelength. Wavelength theoretical values were calculated using our measurements and equation 1, values for observed wavelength in Hibiscus petals were reported in (Lugo et al., 2023), the circles indicate the predicted value using the average values for film thickness and Young’s modulus of the film and substrate, the triangles indicate the predicted values considering the standard deviation for film thickness, and Young’s modulus values. Scale bars: (A) 100 μm, (B) 10 μm, (C) 5 μm.

The results of our AFM experiments to measure the cuticle proper (upper layer) stiffness are also summarized in FIGURE 3E and SUPPLEMENTAL FIGURE S03 for striation onset and open flower stages. These measurements were taken by shallow indentation within the depth range of the cuticle proper, without any dissection or perturbation of the layer structure. The mean Young’s modulus of the cuticle proper at striation onset is around 27.3 MPa, decreasing to around 20.1 MPa at the open flower stage (SUPPLEMENTAL FIGURE S04).

In parallel, we measured the elastic moduli of isolated cuticle thin films through TUTTUT (Liu et al., 2015; Bay et al., 2019). The results of these experiments to analyse the cuticle proper (upper layer) are summarized in FIGURE 3E for striation onset and open flower stages. The mean Young’s modulus of the cuticle proper at striation onset is around 57 MPa, decreasing to approximately 23 MPa at the open flower stage.

All our measurements indicate that the cuticular layer (substrate) is softer than the cuticle proper (upper layer), as hypothesized in previous studies (Airoldi et al., 2021, Huang et al., 2017; Lugo et al., 2023). The difference in the stiffness between these two layers increases throughout time, as the cuticular layer becomes softer during petal growth to a greater degree than does the cuticle proper.

The initial linear slope of the stress–strain response indicates Young’s moduli of the cuticles (FIGURE 3D). We found similar moduli values with both techniques in the open flower, whereas at the onset stage we observe a small difference in the values obtained from the two techniques (FIGURE 3E).

### 3. Wavelength and critical strain in *Hibiscus trionum* petal striation patterns

The striation patterns observed on the *H. trionum* petal cuticle have key features, such as the amplitude (A) and wavelength (λ) (FIGURE 2C). To characterize the pattern wavelength (the distance between two adjacent crests), we used previously published detailed characterization of *H. trionum* cuticle material properties (i.e., cuticle film thickness (h), Lugo et al., 2023) at different stages in combination with the Young’s moduli presented here (TABLE 1). We verified the relationship between stiffness and thickness of our biological materials and wrinkle wavelengths, using Eq. (1) to predict the theoretical wavelength. With the measured moduli and thickness at the onset stage, we calculated a theoretical wrinkling wavelength of 987 nm (FIGURE 3F, TABLE 1), which increases at the mature bud stage (2510 nm), then drops to 2115 nm at the open flower stage. In comparison, the average measured values for the observed wavelength are 1308 nm at the onset stage (ranging from 600 to 2400), 1289 nm at the mature bud (ranging from 500 to 2200), and 1313 nm (ranging from 400 to 2500) in the open flower stage (Lugo et al., 2023). These values were calculated using the average values for cuticle film thickness and Young’s moduli of the cuticle film and substrate (TABLE 1). However, considering the standard deviations of the cuticle thickness (25.7 nm), the Young’s modulus of the cuticle film (9.57) and the Young’s modulus of the substrate (13.48), the predicted wavelength varies from 341 to 1633.54 nm at the striation onset stage (FIGURE 3F).

The theoretical wavelength at striation onset is similar to the observed one, but at the maturity and open flower stages there are notable differences.

## DISCUSSION

Here, we analysed stiffness of a layered living tissue during development to understand the formation of striation patterns in the epidermal surface in *H. trionum* petals. We describe AFM and TUTTUT techniques, which generate reasonably consistent data, providing confidence in the methods applied.

For the use of AFM to calculate Young’s modulus, first, we must assume that the material we measure is isotropic and incompressible. Furthermore, the depth of the indentations and the choice of the indentation shape and size of the cantilever present challenges and can result in errors in the calculations. However, the fact that our AFM Young’s moduli are very similar to the values obtained with TUTTUT (FIGURE 3E) suggests that we can accurately measure petal cuticle stiffness with AFM and provides confidence in our cuticular layer data by extension.

Another important consideration is that measurements of the cuticle proper stiffness using AFM may be affected by the properties of the cuticle layer. Based on the estimated cuticle proper thickness (∼20-60nm) relative to the AFM tip-cuticle contact radius (∼10nm), finite size correction factors (Gao et al, 1992) for stiffer films on softer substrates estimate that errors may be on the order of 10% - 20%. These errors are smaller than other deviations observed in our measurements (FIGURE 3E). Accordingly, rather than focus on absolute moduli values, we measured the stiffness in the cuticular layers over time to calculate the stiffness ratio (cuticle proper/ cuticle layer) that generates the observed striation patterns during petal development in *Hibiscus trionum*. According to our data, the stiffness of the cuticle layers changes over time, both layers become softer at the open flower stage. However, of the two layers it is the cuticle layer for which stiffness decreases most significantly. As a result, the stiffness ratio increases dramatically: at the onset of striation initiation, it is around 3.73 and at the open flower stage it has increased to approximately 18.27. Further analysis will focus on the consequences of changing the timing of cuticle formation and stiffness over developmental time.

The stiffness of cell layers can influence plant development and affect the shape and structure of an organ. Some changes in tissue mechanical properties (including stiffness) can occur during plant development, affecting cell division and expansion. In Arabidopsis roots, differential cell wall stiffness is associated with the elongation or differentiation zones (Liu et al., 2022). In Arabidopsis fruits, the stiffness of the outer layer in the valves is constant during fruit elongation, however, a mutant affected in the degradation of cell wall components displayed an increase in fruit stiffness, affecting fruit elongation (Di Marzo et al., 2021). In the Hibiscus petals, we observed a decrease in stiffness of the cuticle proper over time. This phenomenon could be related to the surface exposure to the environment: the flower bud is closed during development, then opens and is exposed to a different environment with differences in humidity and temperature. Alternatively, given that the flower is open for a relatively short time (few hours), it is possible that the softening represents the onset of senescence, involving changes in cell activity.

With respect to the plant cuticle, elastic moduli have previously been measured in very few systems: in isolated cuticles from fruits (López-Casados 2007), and during tomato fruit growth (Benitez et al., 2021). Analysis on flower petal surfaces has been done in Rose (Almonte et al., 2021). The cuticles in the Hibiscus petals are soft compared to published values for other organs, particularly those of fruit. This big difference in Young’s modulus between cuticles in petals and fruits could be related to the different functions of cuticle in different organs. In fruit the cuticle acts as a barrier to protect against dehydration, pathogens, and insect pests over a much longer time than the lifespan of a petal (Skrzydel et al., 2021).

We observed several changes occurring at the whole petal level, such as increases in cell size and layer thickness. We hypothesize that the stiffness of the cuticular layer could be influenced by growth itself, with cell wall loosening through the activity of proteins such as expansins necessary for cell expansion (Cosgrove, 2015). However, the striation patterns appear to be more stable. As shown in Lugo et al. 2023, the striation wavelength is set when the pattern emerges and remains constant through petal growth. This means that upon striation formation, the deformation seems to be locked despite the changes in cell size we observe from striation onset to flower opening. We hypothesise that this stability of the wrinkling wavelength across petal development could be an effect of the inelastic nature of biological materials during growth. This hypothesis is consistent with our observations after the isolation of the cuticle proper (mechanically or with enzymatic treatment), where the wrinkling patterns were still preserved in the isolated cuticle (FIGURE 3A-C).

When comparing the theoretical versus the observed wavelength values, we observed that the theoretical value is consistent with the range of values measured but not consistent with their median. Our hypothesis is that this discrepancy is generated by changing properties of this living system during development. During the striation onset stage, large parts of the flower are not yet producing striations. Additionally, given the very small surface measured with AFM, we could well be measuring the properties of parts of the tissue that are not yet undergoing wrinkling. Therefore, we are likely to be underestimating the difference in stiffness between the layers during the measurements. This is also suggested by a bigger spread of the AFM data and a bigger discrepancy between the AFM and TUTTUT measurements in the cuticle at the striation onset stage compared with the cuticle of the open flower. Additionally, the wavelength prediction using Eq. (1) only applies to a flat domain i.e., the striation onset. At the later stages, when the cell surface is more curved, we observed bigger differences between the theoretical vs. the observed values. The simulations performed by Lugo et al. (Lugo et al., 2023) have shown how curvature can delay wrinkling onset and therefore could result in a wider wavelength than the one predicted by the formula. The stiffness values of the cuticular layer at the mature bud stage measured with AFM are very similar to the values for the open flower (TABLE 1). However, at the same time the cuticle thickness has increased, and the cuticle stiffness is similar to the values reported for the onset-stage, therefore the theoretical prediction of the wavelength increases to 2.5 μm. This means that during the night that divides the two growth stages (FIGURE 1E) both the growth of the layers and the softening of the substrate will give the right material properties to develop the wavelength observed in the flowers. The dynamic nature of this system and the presence of a curved surface explain the range of wavelengths observed in the flower. It is ultimately this imperfection in the pattern that creates the blue halo that increases salience to pollinators (Moyroud et al., 2017).

The deformations induced by the buckling of the cuticle layers seem to be extremely stable. After we exposed the cuticular layer surface by removing the top cuticle proper, the wrinkle prints were observed (FIGURE 3B,C). Both observations indicate that the strains applied to the materials exceed critical values for the onset of inelastic deformations.

## CONCLUSIONS AND PERSPECTIVES

Here, we report the measurement of stiffness of cuticular layers through petal development. According to our analyses, the AFM and TUTTUT measurements overlap, indicating that both methods could be used to measure stiffness of cuticular layers in *Hibiscus* petals and in other living bilayer systems.

The characterization of *Hibiscus trionum* petals using multiple approaches has given us the possibility to characterize in full the cuticle bilayer in the flower and the buckling phenomenon under the effect of compressive forces. We show that the formula used to describe materials reliably describes our biological system and we can accurately predict the wavelength after buckling by knowing the thickness of the film and the stiffness of the two layers. *Hibiscus trionum* petals, despite being a living and changing biological system, behave and buckle like other materials.

A softer outermost layer would allow a small moduli ratio (R=E_f_/E_s_ >1) and be helpful to achieve the desired small wavelength to enable optical diffraction, and to stabilize a wrinkling pattern (Auguste et al., 2017). However, for plants, a soft cuticle may be insufficient to protect tissue against biological or physical degradation.

Our study also reveals the dynamic development of material properties, with tissue stiffness changing through developmental time. This dynamic property may allow multi-functionality that is difficult to achieve through a static material and may be of interest in the bioinspired design of new materials.

We aim to use the described protocol to dissect the contribution of genetic, biochemical, and environmental factors in the formation of softer or stiffer cuticular layers and their influence in the formation of cuticular patterns. Additionally, further analysis will be conducted to assess the contribution of other cellular features, such as cell curvature and cell shape, to the formation of the observed wrinkling patterns.

## AUTHOR CONTRIBUTIONS

Conceptualization: CAA, CC, AJC, BJG; Methodology: CAA, CC, HF, AJC, BJG; Investigation: CAA, CC, HF; Validation: CAA, CC, HF, CAL; Formal Analysis: CAA, CC, HF, CAL; Resources: AJC, BJG; Writing – Original Draft: CAA, HHU; Writing – Review & Editing: HHU, AJC, BJG; Visualization: CAA, CC, HHU, HF, CAL; Supervision: CAA, CC; Project Administration: AJC, BJG; Funding Acquisition: AJC, BJG.

## ACKNOWLEDGMENTS

We would like to thank Matthew Dorling for excellent plant care and the College of Natural Sciences Greenhouse at UMass Amherst. We are grateful to Alexis Peaucelle for AFM training, and to Alessandra Bonfanti, Colm Durkan and Udhaya Ponraj for useful discussion on the AFM technique and data processing.

We are grateful to the Open Plant Initiative and to the SLCU Microscopy Core Facility, supported by the Gatsby Charitable Foundation. We thank Prof James Watkins for the use of the Zygo profilometer at UMass Amherst. This work was supported by a HFSP grant RGP0019/2017, a BBSRC grant BB/P001157/1 and the Cambridge University Botanic Garden Research Fund.

---

**Supplemental figure S01.**
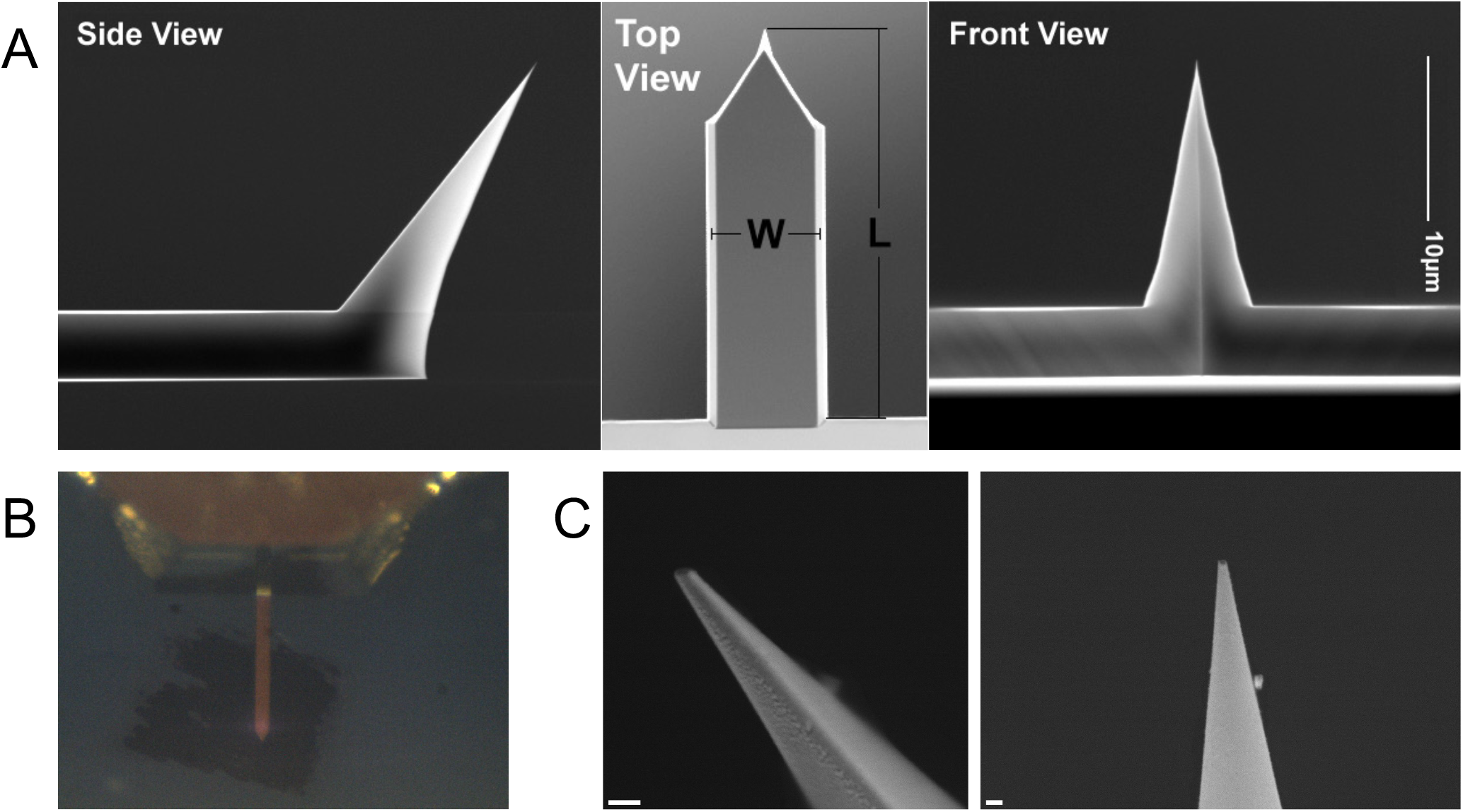
Overview of the tip used in the atomic force microscopy (AFM) experiments. A) Side, top, and front view of the tip. Images provided by the manufacturer. B) Top view of the tip taken with the AFM built-in camera or C) with the scanning electron microscope. Scale bars in C represent 200 nm.

**Supplemental figure S02.**
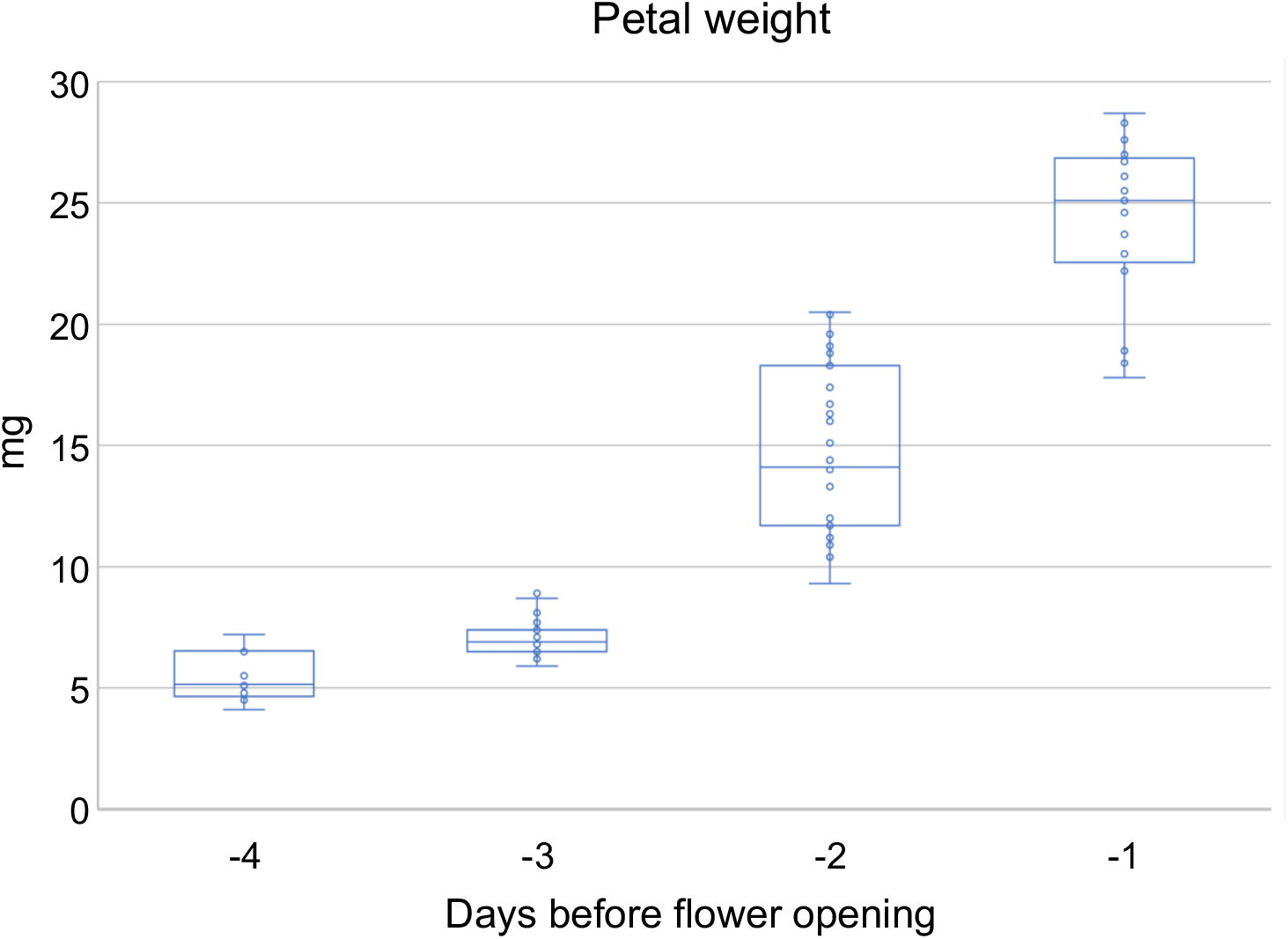
*Hibiscus trionum* petal mass increase over 8me. Weight of single petals from flowers before opening. Bud size was used to es3mate 3me before opening.

**Supplemental figure S03.**
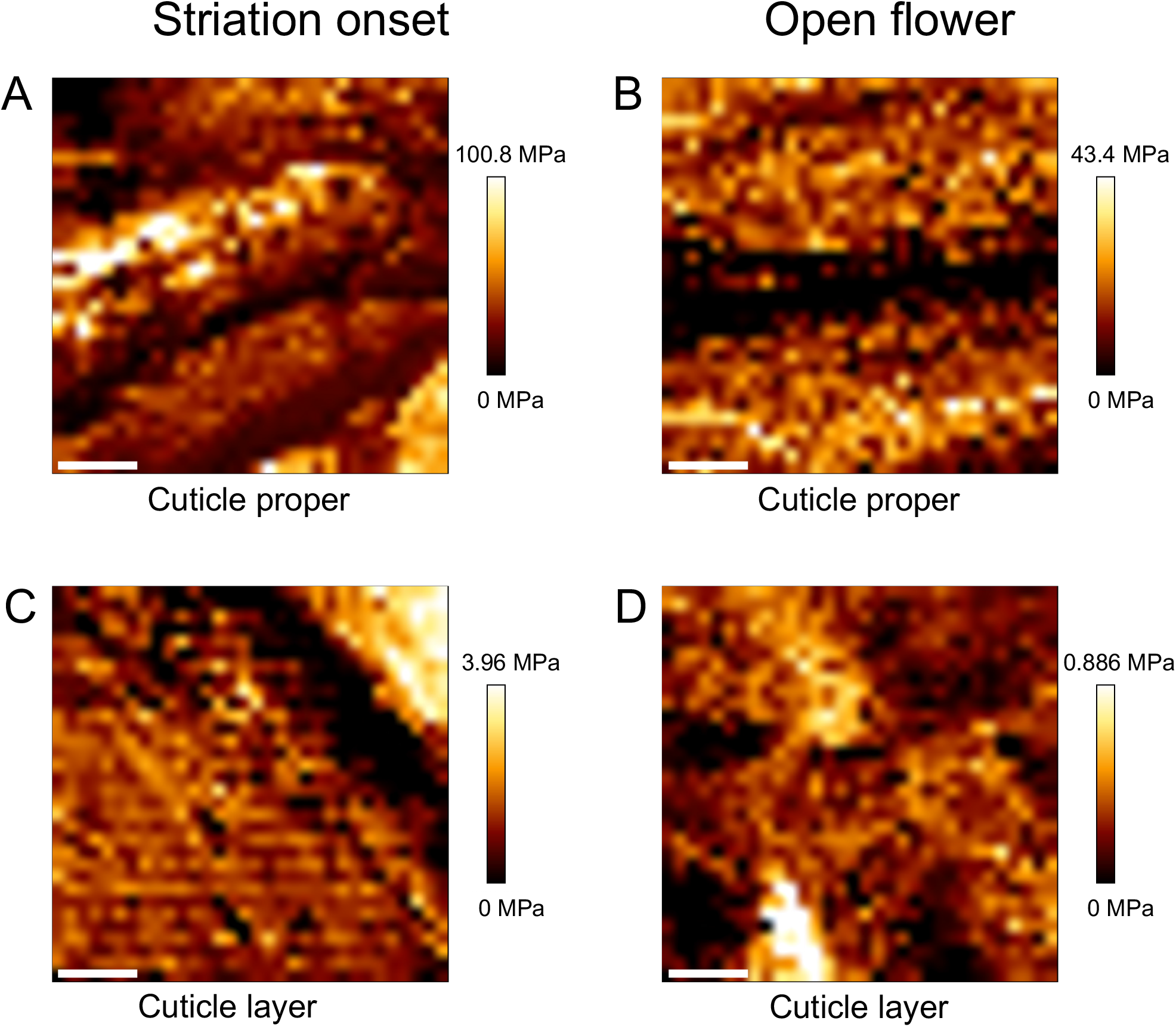
Atomic force microscopy (AFM) maps showing Young’s modulus for the cuticle proper and cuticle layer at the stria3on onset and at the open flower stage. Scale bars represent 200 nm.

**Supplemental Figure S04.**
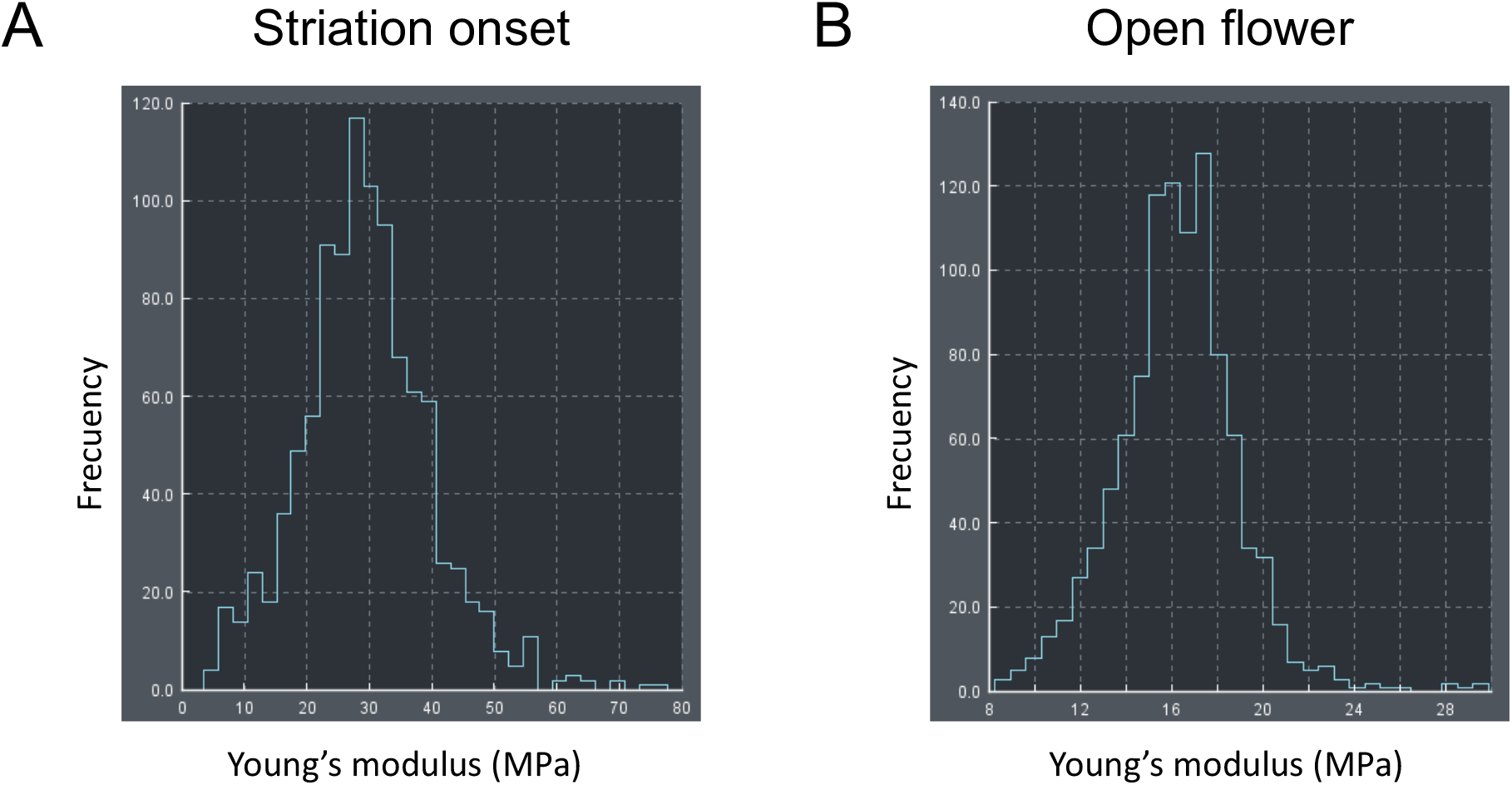
Comparison of Young’s modulus values in the cu8cle proper in *H. trionum* petals. One example of the Young’s modulus values measured with Atomic Force Microscopy at the stria3on onset and at the open flower stage.

**Supplemental figure S05.**
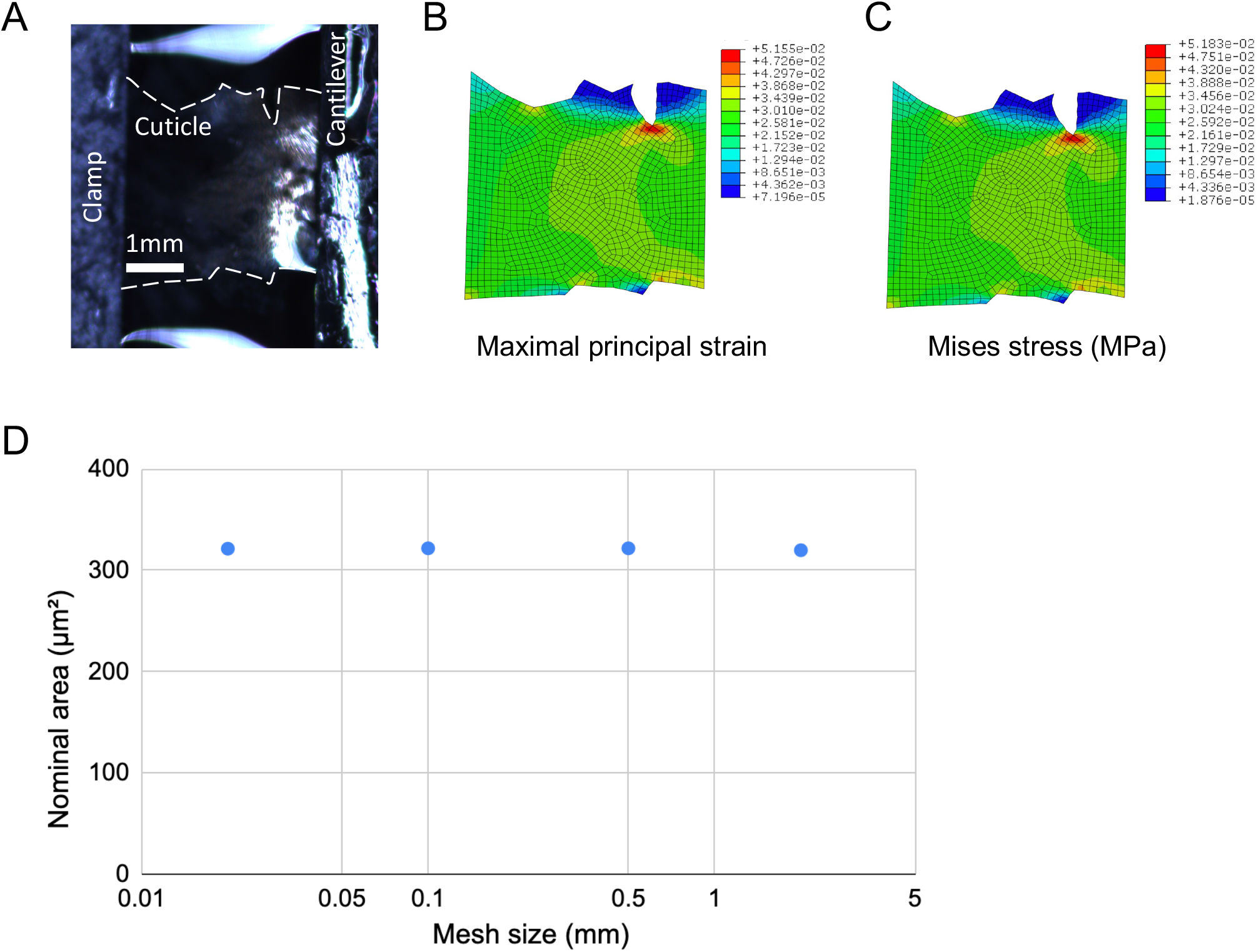
Finite element calibration of TUTTUT experiment. A) TUTTUT experiment on a petal cuticle in the striation onset stage. The floated cuticle layer was adhered to a cantilever at right side and a loading clamp at leP side, subject to a tension in horizontal direction. Linear elastic finite element calculation on a cuticle model with the same shape and thickness. B) The maximal principal strain distribution and C) the von Mises stress distribution with an overall strain of 2 %. (D) The mesh sensitivity study with mesh from 0.02 mm to 2 mm. The calculated nominal area has 0.57 % variation.

